# Phosphoregulation provides functional specificity to biomolecular condensates in the cell cycle and cell polarity

**DOI:** 10.1101/796755

**Authors:** Therese M. Gerbich, Amy S. Gladfelter

**Affiliations:** Department of Biological Sciences, University of North Carolina at Chapel Hill, Chapel Hill NC 27599; Marine Biological Laboratory, Woods Hole, MA, 02543

## Abstract

Cytoplasmic patterning is a feature of many cells from embryos to neurons and fungi. Biomolecular condensation is a way of organizing cytosol in which proteins and nucleic acids coassemble into compartments. How molecular identity of condensates is achieved is not well understood. In the multinucleate, filamentous fungus *Ashbya gossypii*, the RNA-binding protein Whi3 regulates the cell cycle and cell polarity through forming macromolecular structures that behave like condensates. Whi3 has distinct spatial localizations and mRNA targets making it a powerful model for how, when and where specific identities are established for condensates. Using mass-spectrometry, we identified residues on Whi3 that are differentially phosphorylated under specific conditions and generated mutants which ablate this regulation. This yielded separation of function alleles that were functional for either cell polarity or nuclear cycling but not both. This study shows that phosphorylation of individual residues on molecules in biomolecular condensates can provide specificity that give rise to distinct functional identities in the same cell.

**Summary:** Residue specific phosphorylation of the RNA-binding protein Whi3 is used to specifically regulate subsets of functionally distinct condensates in the multinucleate fungus *Ashbya gossypii*.

## Introduction

A central feature of cell organization is the assembly and regulation of compartments. Cellular biochemistry is spatially limited by sequestration into both membrane-bound organelles and membraneless bodies, frequently termed biomolecular condensates (Banani et al., 2017). While it is increasingly clear that there are many condensates in both the cytosol and nucleus, the mechanisms that control where and when they form, determine their physical properties and distinct functions remain under intensive study (Brangwynne et al., 2011; Banani et al., 2017). Physical principles for phase transitions indicate that condensates should be sensitive to local changes in the abundance of key components, changes in the solvent conditions or electrostatic interactions amongst components.

Work on the RNA-binding protein Whi3 in the filamentous fungus *Ashbya gossypii* has led to some key insights into how specificity can arise. This protein forms assemblies dependent on an extended polyQ-tract and binding to specific mRNAs. *CLN3* (a G1 cyclin) transcripts are heterogeneously positioned near nuclei in a Whi3-dependent manner. *BNI1* (a formin) transcripts are positioned at growing tips and nascent polarity sites in a Whi3-dependent manner. The assembly of these Whi3 structures is essential for asynchronous division of nuclei in a common cytoplasm and the generation of new polarity sites (Lee et al., 2013; Lee et al., 2015). The fact that this single protein forms functionally distinct condensates in a shared cytoplasm makes it an excellent model to examine sources of specificity. Our previous work identified a key role for RNA secondary structures in promoting molecular specificity and specifically the capacity of mRNAs involved in the cell cycle and cell polarity to engage in RNA-RNA interactions (Langdon et al., 2018). However, while RNA is very important for maintaining complex identity, it is not well understood what cues drive the assembly and disassembly of these complexes.

We hypothesized that localized signaling may generate specific post-translational modifications on Whi3 that could control where and when it condenses, trigger dissembly, or direct the function of a given complex. *Ashbya* can easily grow to be hundreds of microns across with dozens of simultaneously elongating tips and nuclei that often travel long distances from their birthplace (Gladfelter et al., 2006; Schmitz et al., 2006). Given their large size and the potential to experience different environments in different parts of the cell, it could be advantageous for cells to use post-translational modifications controlled by local signaling related to the cell cycle or polarity. Cell cycle and nutrient signaling are well known to involve kinase signaling (Rhind and Russell 2012; Conrad et al., 2014), and phosphorylation could regulate the components or activities of a condensate. If the modifications are stable, they could lead to condensate structures that persist over long timescales and lengthscales. If condensates diffuse or are transported, they may fuse with other Whi3 assemblies in a new region of cytoplasm to share information, and still be able to be rapidly disassembled in response to a new local cue. In this way, PTMs may be a way for locally generated signals to be shared over a long length and timescale for the cell to integrate information and/or to create a spatially-restricted functional condensates.

The genetic tractability and clear phenotypic links to function for the Whi3 protein provide an opportunity to search for specificity at the level of post-translational modification of a protein that is a common component of different condensate assemblies. Remarkably, we identified clean separation of function mutants on two key sites on the protein that specifically block either the nuclear cycle or cell polarity functions. This indicates that despite seemingly low complexity sequences that there is exquisite sensitivity of condensates to even small changes in local charge.

## Results and Discussion

### Identification of phosphorylation sites on Whi3

We hypothesized that the assembly and/or dissolution of Whi3 condensates is controlled at specific places and particular times in the nuclear division cycle. In support of this idea, when cells are arrested in nocodazole which leads to a G2/M arrest, there is a substantial decrease in the number of Whi3 puncta in the vicinity of nuclei (Fig. 1A&D). This suggests that the assemblies are dissolving prior to or in mitosis. In acute heat shocked cells (2h at 42°C) additional puncta formed in the hyphae, suggesting a redistribution of the protein into potentially distinct stress-induced condensates (Fig 1A&D). These conditions did not affect the number of hyphae containing Whi3 puncta at their tips (Fig. 1C), and overall protein levels remained similar (less than two-fold change in either condition) (Fig. 1B). We hypothesized that rapid changes in the state of condensates could be triggered by post-translational modifications to Whi3 protein.

**Figure 1.**
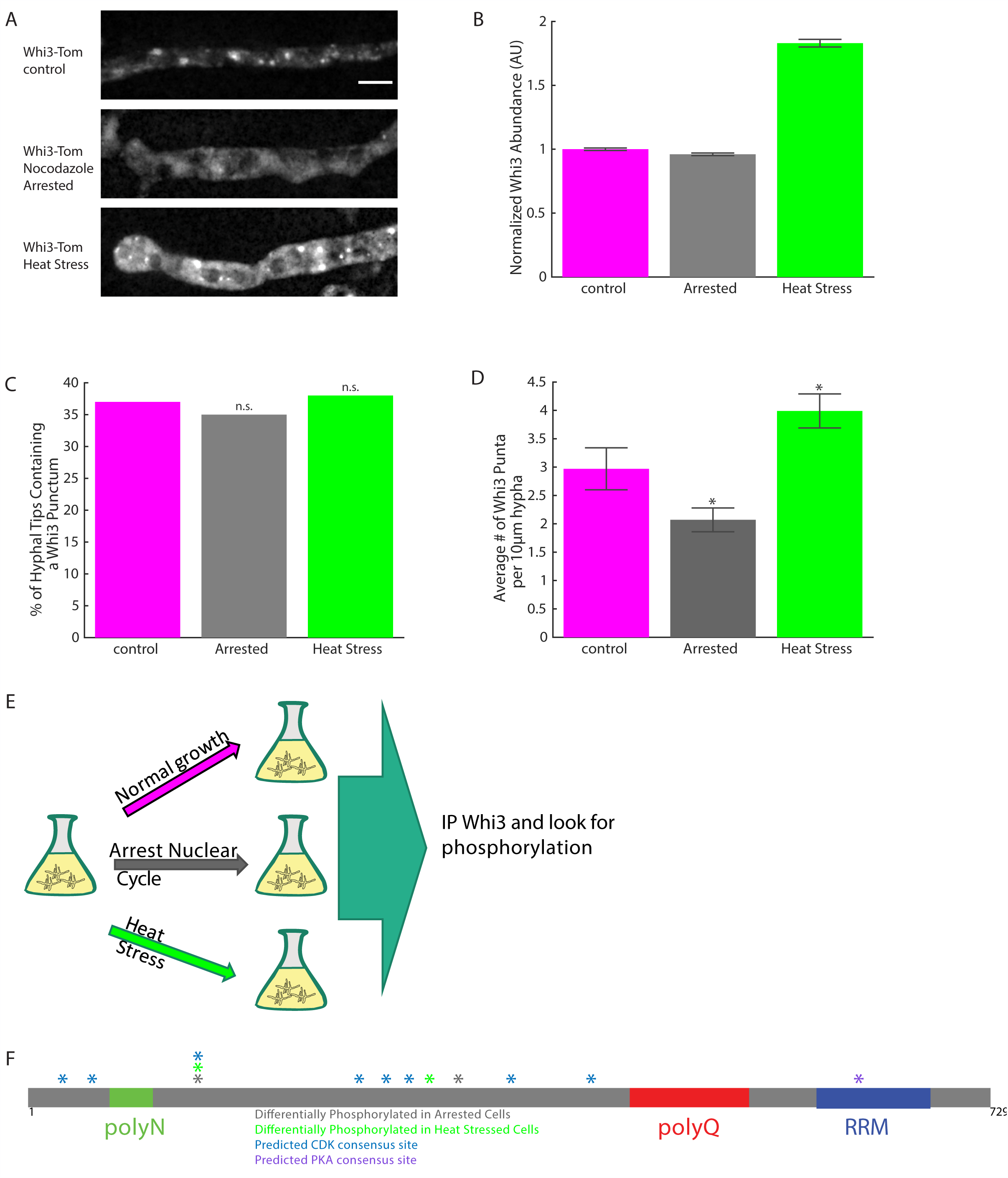
Whi3 is differentially phosphorylated during nocodazole arrest and heat stress. (A) Representative images of Whi3-tomato protein localized in hyphae under different growth conditions. Scale bar 5 μm. (B) Normalized Whi3 abundance as measured by fluorescence intensity of Whi3-tomato protein within the hypha under different growth conditions. Bars denote SE. (C) Percent of hyphal tips containing a Whi3 punctum. \**p* < 0.05 by N-1 Chi-Square test. (D) Average number of Whi3 puncta per 10 μm hyphal segment. Bars denote SE. \**p* < 0.05 by T-test. (E) Schematic of mass spectrometry workflow. (F) Diagram of sites that were more phosphorylated during nocodazole arrest and/or heat stress. Also included are predicted CDK and PKA sites.

We therefore evaluated if Whi3 was differentially phosphorylated under cell cycle arrest or heat stress. Whi3 was immunoprecipitated from *Ashbya* cells growing asynchronously, arrested in mitosis with nocodazole treatment or under heat stress. The immunoprecipitate was analyzed by mass spectrometry to identify phosphorylation sites on Whi3 protein in each of the different conditions (Fig. 1E). We found multiple (X=22, Table 1) of sites on the protein that were phosphorylated and several sites on the protein that were more phosphorylated in nuclear cycle arrest and heat stress conditions. The mass spectrometry did not fully cover the entire Whi3 protein and notably repetitive sequences in the polyQ region as well as portions of the RNA-binding domain were not captured. We then replaced Whi3 at the endogenous locus under the native promoter with non-phosphorylatable (S>A) and phosphomimetic (S>D) versions of the protein for each of the suspected sites of regulation generated by the mass spectrometry results as well as well as predicted CDK and PKA sites (Mok et al., 2010) (Fig. 1F). Alleles were tagged with tdTomato at the C-terminus to analyze the impact of mutations on Whi3 puncta distribution. We then assayed these mutants for cell behaviors regulated by Whi3 condensation: nuclear asynchrony and polarized growth and then evaluated the localization of the mutant protein.

**Table 1.** Sites phosphorylated with any confidence.

|  |  | Aynchronous<br>1 | Arrested<br>1 | Heat<br>Stress<br>1 | Asynchronous<br>2 | Arrested<br>2 | Heat Stress<br>2 |
| --- | --- | --- | --- | --- | --- | --- | --- |
| S025 |  | C | ? | NC | C | NC | C |
| S026 |  | X | X | NC | ? | NC | X |
| T034 | CDK-1 | X | X | NC | C | C | C |
| S039 | CDK-2 | C | X | NC | C | C | X |
| S111 | CDK-3 | X | X | NC | X | X | X |
| S198 | GIN4-3 | X | NC | NC | C | C | C |
| S249 | CDK-4 | X | X | NC | X | X | X |
| S283 | GIN4-4 | X | X | X | X | C | X |
| S289 |  | C | X | NC | C | X | C |
| S292 |  | ? | ? | NC | X | C | X |
| T294 | YAK1-1 | X | C | NC | ? | C | C |
| T313 |  | ? | ? | C | C | C | X |
| T314 | CDK-6 | ? | ? | C | ? | ? | ? |
| S355 |  | X | X | NC | X | X | X |
| T359 | CDK-7,<br>HOG1-1 | X | X | NC | C | C | C |
| S372 | GIN4-5,<br>PKA-1 | ? | ? | NC | C | C | X |
| S373 | GIN4-6 | NC | C | NC | C | C | ? |
| S396 |  | NC | NC | NC | ? | ? | C |
| S413 |  | ? | X | X | C | NC | C |
| S415 |  | C | C | C | NC | C | ? |
| S559 |  | NC | X | NC | NC | C | X |
| S561 |  | ? | C | NC | NC | C | C |
| S583 |  | NC | X | NC | X | C | ? |

Coverage of Predicted Kinase Sites not found
|  |  | Aynchronous<br>1 | Arrested<br>1 | Heat<br>Stress<br>1 | Asynchronous<br>2 | Arrested<br>2 | Heat<br>Stress<br>2 |
| --- | --- | --- | --- | --- | --- | --- | --- |
| S024 | GIN4-1 | C | C | NC | C | C | C |
| **T183** | GIN4-2 | NC | NC | NC | C | C | C |
| S271 | CDK-5 | C | C | NC | C | C | C |
| S373 | GIN4-6 | C | C | NC | C | C | ? |
| T420 | CDK-8 | NC | C | NC | C | C | C |
| S556 | GIN4-7 | NC | C | NC | NC | NC | C |
| **S637** | GIN4-8,<br>PKA-2 | NC | NC | NC | NC | NC | NC |

C= region covered by mass spec
NC= region not covered by mass spec

We found a spectrum of phenotypes that could be assigned into 4 distinct classes: complete loss of function with defects in both nuclear cycling and polarity, partial loss of function in nuclear cycling or polarity, or no phenotype (Table 2, Figs. 2-4). Importantly, all mutants are comparably expressed to wild-type protein indicating that the defects are not due to mis-expression of the mutant alleles (Fig.4C). None of the mutant alleles substantially changed the distribution of nuclei in different cell cycle phases indicating that the cell cycle defects primarily impacted synchrony rather than progression through the division cycle per se (Fig. 2D). In some cases, phenotypes were associated with the loss of detectable Whi3 condensates but in other cases condensates appeared morphologically normal and abundant but were clearly not functional (Fig. 4). Below, we discuss the most interesting cases of clear separation of function alleles in detail and then examine the associations between localization and function.

**Table 2.** Summary of Cellular Phenotypes and Whi3 Localization.

|  | **Polarity** | **Tip**<br>**Puncta** | **Asynchrony** | **Hyphal**<br>**Puncta** |
| --- | --- | --- | --- | --- |
| **ctrl** | + | + | + | + |
| **111A** | + | + | + | + |
| **111D** | + | - | - | -- |
| **283A** | + | + | + | + |
| **283D** | + | + | + | + |
| **289A** | - | ++ | + | + |
| **289D** | -- | ++ | - | + |
| **637A** | -- | -- | + | - |
| **637D** | -- | -- | - | -- |
| **cdkA** | - | ++ | - | ++ |
| **cdkD** | -- | ++ | - | - |

**Figure 2.**
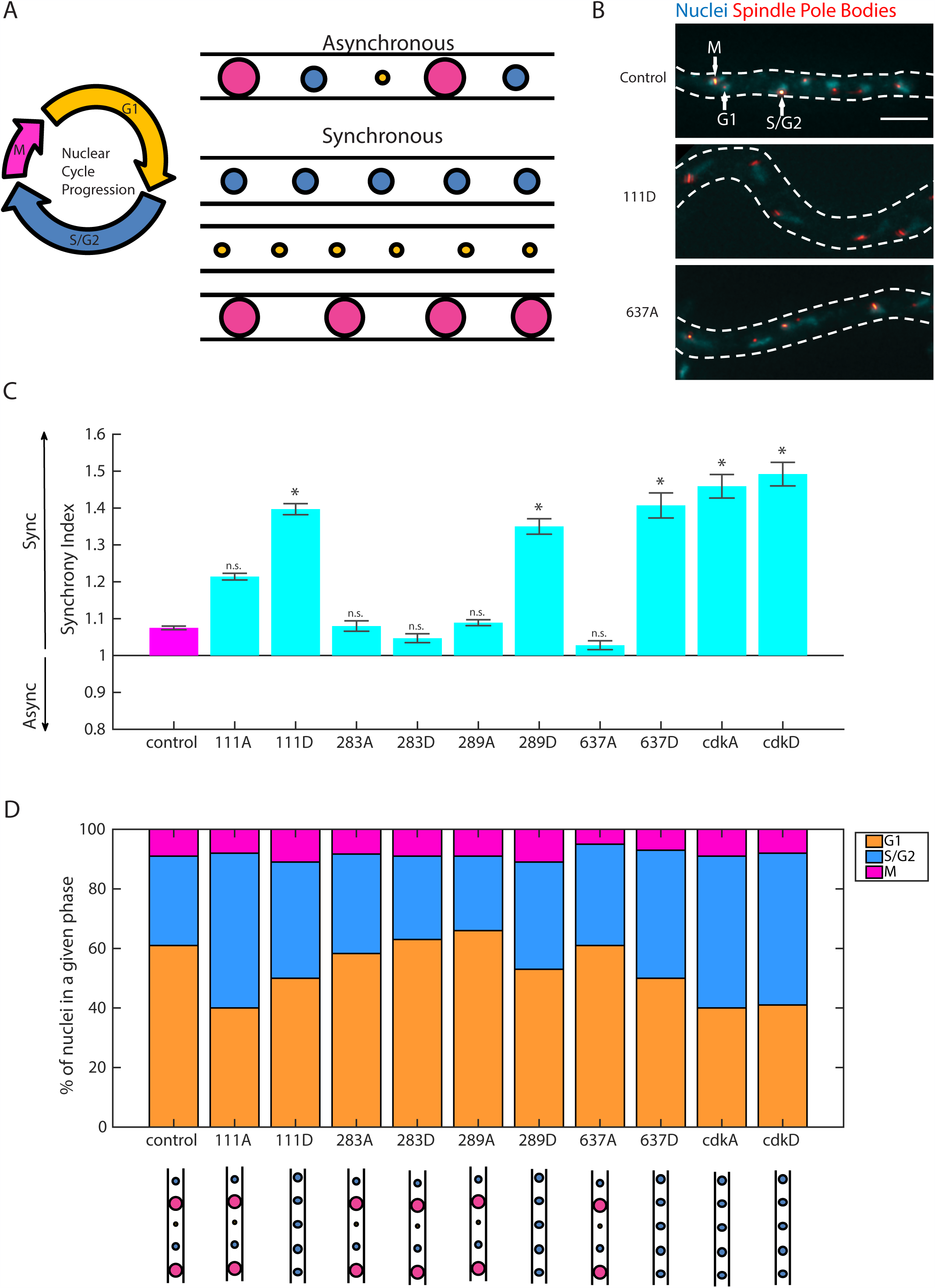
Single point mutants at sites of Whi3 phosphorylation can increase nuclear synchrony. (A) Nuclei in wild type *Ashbya* cells go through the cell cycle asynchronously, but can become more synchronous under certain genetic or environmental conditions, including removal of the polyQ tract from Whi3. (B) Representative images of SPB labeling for synchrony scoring in control (asynchronous), 111D (increased synchrony mutant), and 637A (asynchronous mutant). SPBs are in red and nuclei are in cyan. Hyphal outlines are dotted lines and representative nuclei of each nuclear cycle stage are indicated with arrows in control cell. Images are maximum intensity projections. Scale bar 5 μm. (C) Synchrony indices for Whi3 mutants as compared to control cells. Bars denote SE. (D) Percent of nuclei in each phase of the nuclear cycle in control and Whi3 mutant strains. G1 nuclei are orange bars, S/G2 nuclei are blue bars, M nuclei are pink bars. Synchrony phenotype for each strain is represented by cartoon image below strain name.

**Figure 3.**
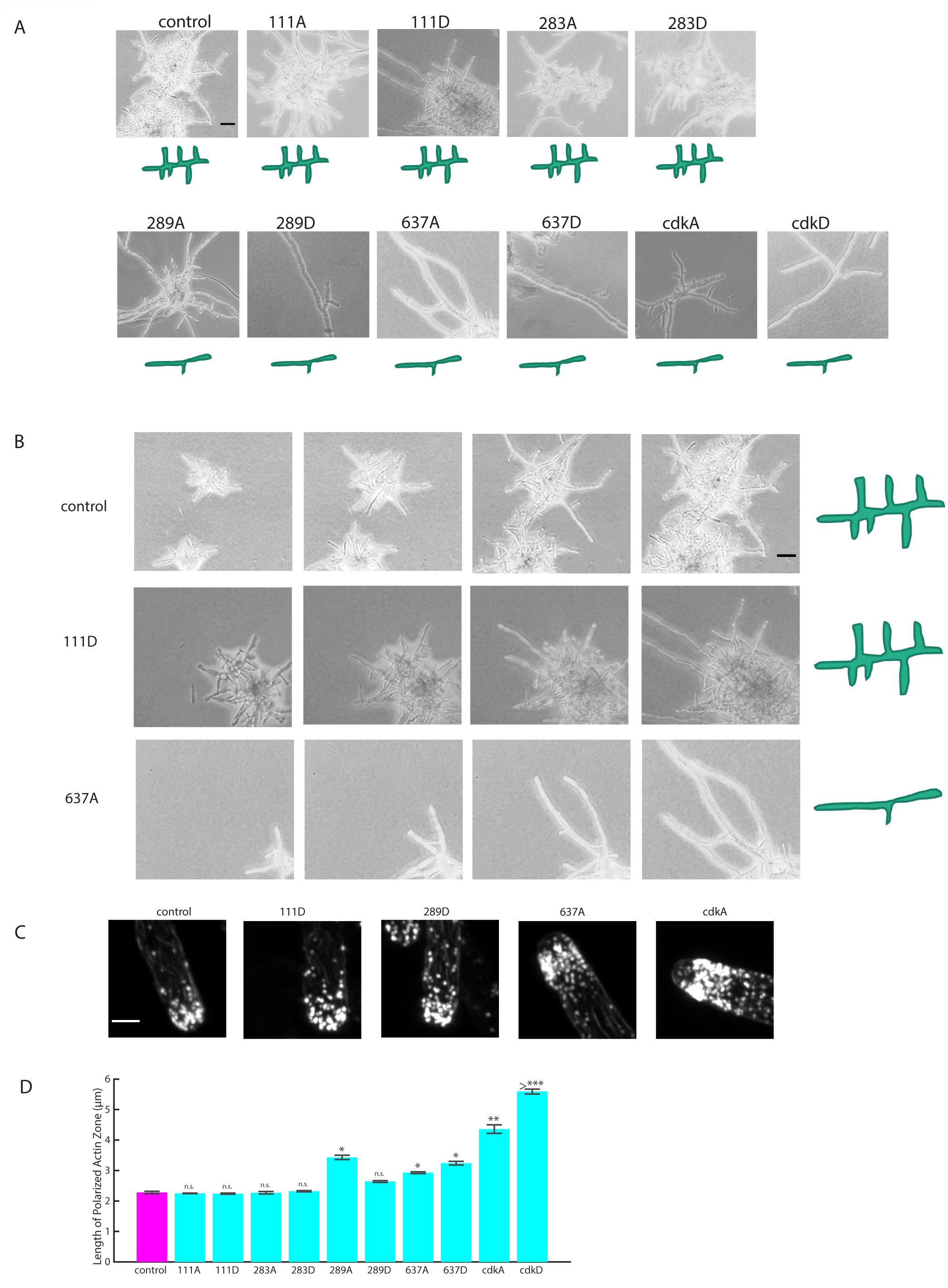
Single point mutants at potential sites of Whi3 phosphorylation can disrupt cellular polarity and lateral branching. (A) Representative images to show gross morphology of control and Whi3 mutant strains grown on agar pads. Cartoon representation of polarity phenotype for each strain is shown below each strain. Scale bar 50 μm. (B) Montage of cells growing on plate for control (normal polarity), 111D (normal polarity mutant), and 637A (disrupted polarity mutant). Interval between frames is 108 minutes. Scale bar 50 μm. (C) Representative images of actin at hyphal tips in control and selected Whi3 mutant strains. Images are maximum intensity projections. Scale bar 2 μm. (D) Length of polarized actin zone along hyphal long axis in control and Whi3 mutant strains. Bars denote SE. \**p* < 0.05 by T-test.

**Figure 4.**
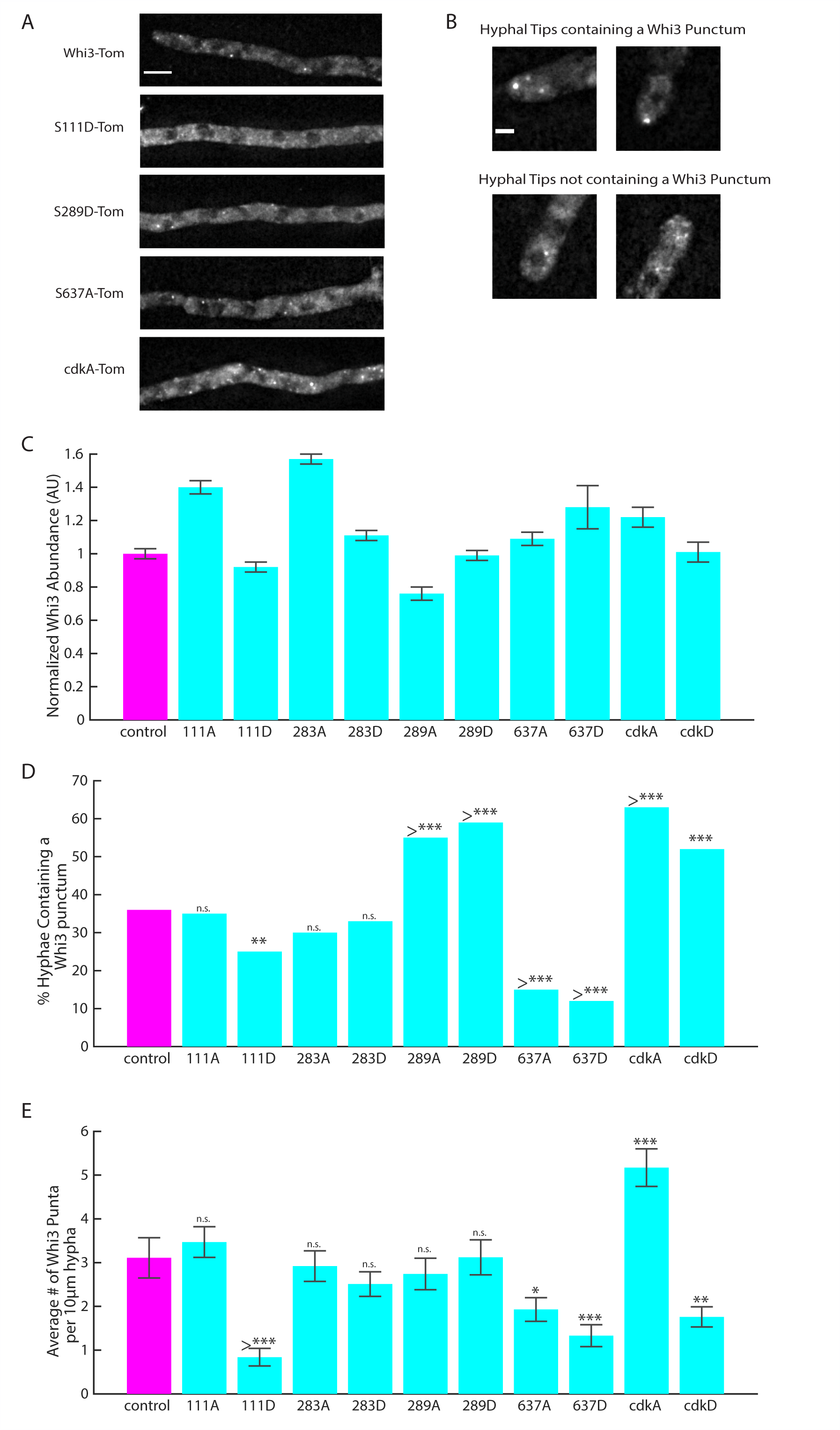
Single point mutants at potential sites of Whi3 phosphorylation can strongly influence Whi3 localization and assembly. (A) Representative images of Whi3 protein in control or Whi3 mutant strains localized in the hypha. Images are maximum intensity projections. Scale bar 5 μm. (B) Representative images of hyphal tips containing or not containing Whi3-tomato puncta. Images are maximum intensity projections. Scale bar 2 μm. (C) Normalized Whi3 abundance in control and Whi3 mutant cells under different growth conditions. Bars denote SE. (C) Percent of hyphal tips in control and Whi3 mutant strains containing a Whi3 punctum. \**p* < 0.05 by N-1 Chi-Square test. (E) Average number of Whi3 puncta per 10 μm hyphal segment in control and Whi3 mutant strains. Bars denote SE. \**p* < 0.05 by T-test.

### Separation of function mutants specific for asynchrony or cell polarity

A major goal of this study was to determine if PTMs could regulate Whi3 condensates in time and space and promote their functional specificity. Two clear separation of function alleles emerged from this analysis supporting that phosphoregulation is a key cue for functional specialization. One partial loss of function allele specifically impacted nuclear asynchrony; cells expressing S111D mutations are significantly more synchronous than wild-type cells (Fig. 2C) but show normal branching patterns (Fig. 3A&B). S111D is a predicted Cyclin-dependent kinase (CDK) site that we detected as a phosphorylation event more frequently in both nocodozole synchronized cells as well as in heat stressed cells. Consistent with the loss of asynchrony phenotype, S111D mutants showed fewer Whi3 puncta associated with nuclei (Fig. 4 E). There is also a modest, but statistically significant, decrease in tip-associated puncta but presumably still enough assembly to support normal polarity patterning (Fig. 3, 4D), and no difference in the length of the polarized actin zone in these cells compared to controls (Fig. 3D). Thus, addition of negative charge at this single residue is sufficient to specifically impact the assembly and function of Whi3 in the cell cycle without impacting functions in cell polarity.

The second separation of function allele we identified was S637A. This site is located within the RRM of the protein and is a conserved, putative PKA consensus site that has been shown experimentally to be phosphorylated in *Saccharomyces cerevisiae* (Mizunuma et al., 2013). The S637A mutant has normal nuclear asynchrony (Fig. 2C) but grossly disrupted polarity (Fig. 3 A&B) with very limited initiation of lateral branches indicating that Whi3 has specifically lost its ability to regulate polarity without impacting nuclear cycling. There is a 50% reduction in tips with Whi3 puncta compared to wild-type cells in the S637A allele which is consistent with the substantial polarity problems (Fig. 3, 4) and an increase in length of the polarized actin zone at tips (Fig 3D). There are also somewhat fewer puncta in the cytosol away from tips (Fig. 4E) which may be a specific loss of the subset of polarity-involved puncta used at the initiation of lateral branches.

Thus, separation of function alleles were identified for both the key processes controlled by Whi3 condensates in *Ashbya*. Remarkably, these single point mutations were each able to recapitulate what was seen in the full polyQ deletion mutant for each process (Lee et al., 2013; Lee et al., 2015). But unlike the polyQ mutants, in both cases, only a specific subset of Whi3 assemblies was lost and the corresponding cellular function was impaired. This indicates that while there may still be regulation that impacts all Whi3 assemblies, each subset can also be controlled independently and it is possible to genetically separate these functions of Whi3.

There has been an emerging understanding of how post-translational modification can be used as a cue to regulate assembly or disassembly of condensate structures in other systems (Wippich et al.,2013; Hofweber and Dormann, 2018). To our knowledge, however, it has not been shown how post-translational modifications can be used to specify different functions for condensates that share common components. Our results indicate that beyond the emergent properties of weak, multivalent interactions, there are likely to be specific and switchable interactions that are important for assembly. Post-translational modifications may lead to different client proteins which can be recruited that impart functional specificity to condensates that are relevant in early stages of assembly. It is remarkable that even single residue changes are sufficient to block assembly and function of specific complexes, indicating that even in the process of phase separation that is seemingly buffered by multivalency that there is critical sensitivity in certain sites of the constituents.

### Whi3 residues phosphorylated during heat stress do not affect polarity or nuclear cycle regulation

The phosphosite screen also identified one site, S283, that was specifically targeted during heat stress. Mutations at site S283 did not have any effect on nuclear synchrony or polarized growth (Fig. 2&3). Similarly, there was no difference in the number of hyphal tips containing a Whi3 punctum between wild type and either of these mutants (Fig. 4D), consistent with the fact that there was no polarity defect in these strains. With respect to hyphal Whi3 puncta, both S283 mutants were indistinguishable from wild type, consistent with them having no problem maintaining nuclear asynchrony (Fig. 4E).

In *Saccharomyces cerevisiae*, Whi3 has been seen to localize to stress granules and P bodies following certain stress conditions (Cai et al., 2013; Mizunuma et al., 2013). We hypothesize that a subset of hyphal Whi3 puncta may be used to regulate other Whi3 phenotypes related to environmental stress response, and S283 might be involved in the regulation of assemblies not seen under normal conditions. When S283 cells were imaged following heat stress, we saw substantially more Whi3 puncta in the S283A cells compared to control (Fig. S1), consistent with this residue playing a role in regulating Whi3 assemblies involved in a heat stress response. In S238D cells we also observed a modest decrease in the number of Whi3 puncta following heat stress (Fig. S1), raising the possibility that phosphorylation at S283 during environmental stress might be used to disassemble existing Whi3 assemblies or even to prevent aberrant or excessive Whi3 recruitment to stress granules.

### Putative CDK-target residues are important for Whi3 function

At the onset of this project, we were concerned that single residues may be insufficient to generate a phenotype so in parallel we made mutants where eight predicted CDK sites were mutated to be phosphomimetic (Ser/Thr to Ala) or non-phosphorylatable (to Asp). Of these eight sites, five of them were found to be phosphorylated in one or more conditions in our mass spec, while the other three sites were not detected in our conditions. Both the cdkA and the cdkD mutant showed extreme defects in both polarity and nuclear asynchrony (Fig. 2 and 3). Surprisingly, both the strains showed an increase in the amount of Whi3 puncta at hyphal tips despite significant defects in generating lateral branches. This supports a role for CDK/cyclin in cell polarity, which would be predicted based on the localization of Cln1/2 cyclin at tips (Gladfelter et al., 2006).

Interestingly, even though both strains showed pronounced defects in nuclear synchrony, the cdkA strains had increased Whi3 hyphal puncta while the cdkD strains had very few. These changes in the frequency of inter-region puncta suggest that phosphoregulation is an important cue for the assembly and dissolution of Whi3 puncta. However, when combined, the localization data also indicate that simply the presence of the puncta is insufficient for normal activity in controlling the division cycle.

### Alleles with loss of function in polarity and nuclear asynchrony

In addition to the CDK mutants, there were two other alleles that showed complete loss of function in asynchrony and cell polarity: S289D and S637D. Interestingly, while the S637A strain does show separation of Whi3 function, the corresponding phosphomimetic S637D strain has both aberrant polarity and a more synchronous nuclear cycle, which cannot be easily explained by the site being used exclusively to regulate Whi3 for its function in polarized growth. The S637D cells have also lost both tip and hyphal puncta. There are a number of potential interpretations of what might actually be going on, but one possibility is that phosphorylation of S637 is required for polarized growth but not for nuclear asynchrony. However, the S637D mutation is not an appropriate mimic for a true phosphorylation event and has created a misfunctional protein that can no longer fulfil any of its normal functions.

In the case of the S289, the phosphomimetic mutants are highly synchronous (Fig. 2C) and show substantial polarity defects (Fig 3A). Notably, these phenotypes are not associated with Whi3’s inability to form puncta as in fact there are still hyphal puncta and even more tips with polarity associated puncta in this mutant (Fig. 4 D&E). We hypothesize that regulated phosphorylation at S289 is important for recruitment of other factors that are needed to make Whi3 complexes fully functional for cell polarity and nuclear asynchrony.

### Loss of function not always associated with loss of condensation

Previous work has shown that when the polyQ tract is removed from Whi3 (ΔpolyQ), the protein is found far less often in puncta (both hyphal and at tips), because its ability to form assemblies has been impaired. In vitro this is seen as an inability to undergo a liquid-liquid phase separation with target RNAs (Lee et al., 2013; Lee et al., 2015; Zhang et al., 2015). The situation is far more complex with the phosphomutants where a substantial fraction of the mutants still form higher-order assemblies in cells and in some cases are even more abundant than the condensates made by wild-type protein. As noted above, both the S289 and CDK alleles all form abundant tip puncta but cannot branch normally and the S289D and cdkA mutants show normal to abundant hyphal puncta but a nuclear synchrony phenotype. This raises the possibility that the purpose of forming these structures is not simply to localize RNA. These alleles demonstrate an important role for phosphoregulation beyond controlling the assembly state but also in the function potentially thought mediating specific protein or RNA interactions.

### Conclusion

In conclusion, this study shows that phosphorylation of a single residue can specifically regulate a subset of condensate assemblies built from a common scaffold protein. Furthermore, in addition to stabilizing or destabilizing the assemblies themselves, phosphorylation can also be used to regulate the activity of assemblies, possibly through the recruitment of other client proteins. Overall, this work advanced our understanding of the molecular grammar used to regulate condensates in the cell.

## Acknowledgements

We would like to thank the Gladfelter and Lew Labs for useful discussions. The work was supported by NIH R01-GM-081506. The authors declare no competing financial interests.

## Author Contributions

T.M. Gerbich conducted experiments. T.M. Gerbich and A.S. Gladfelter designed experiments and wrote manuscript.

## Materials and Methods

### Strain Construction

To create Whi3 mutant strains tagged with tomato, geneblocks containing the mutations of interest were ordered from IDT (Table 3). These geneblocks were then used to replace the relevant portions of Whi3 in AGB993 using Gibson cloning to amplify the geneblocks and relevant portions of the plasmid (Table 4). Constructs were sequenced across the Whi3 region of the plasmid to ensure that no additional mutations had accrued during the cloning process (Table 5). Each plasmid was then cut by SacII, XhoI, and NdeI. The 6.377 kb fragments containing the mutant Whi3, the fluorescent tag, and the NAT resistance marker were separated and purified using an agarose gel and integrated into *Ashbya* using protocols described previously (Wendland et al., 2009) (Table 6).

**Table 3.**
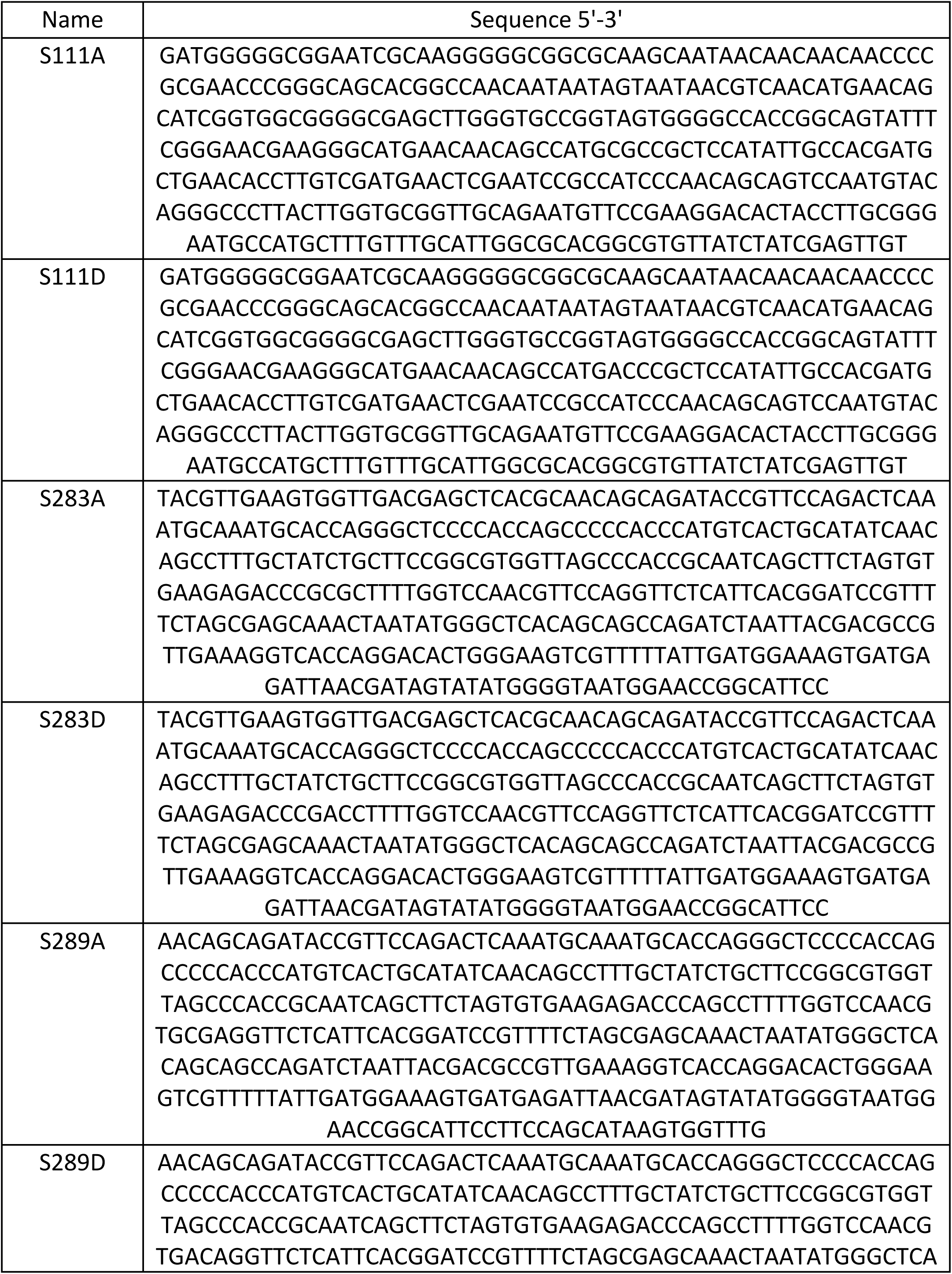
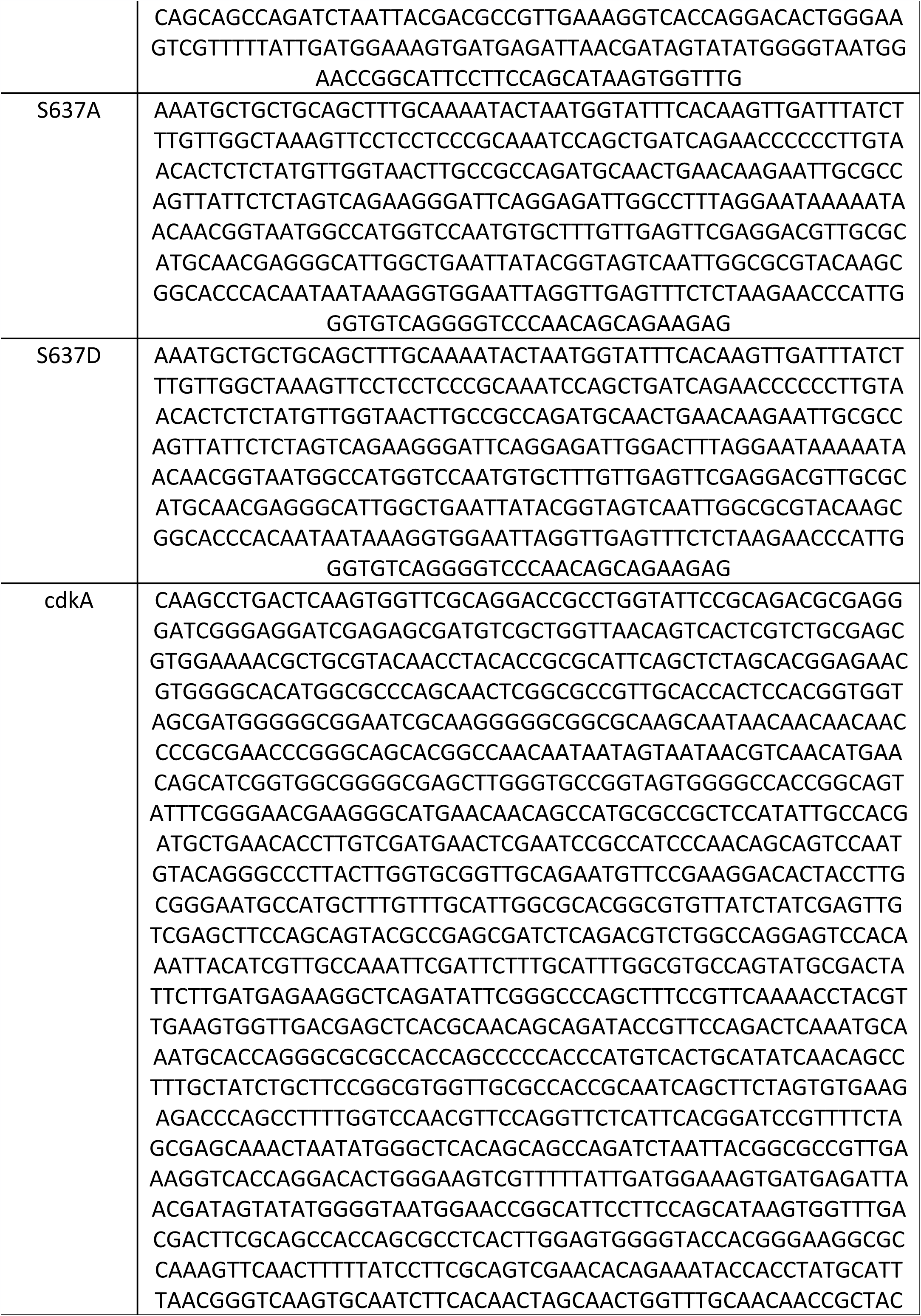
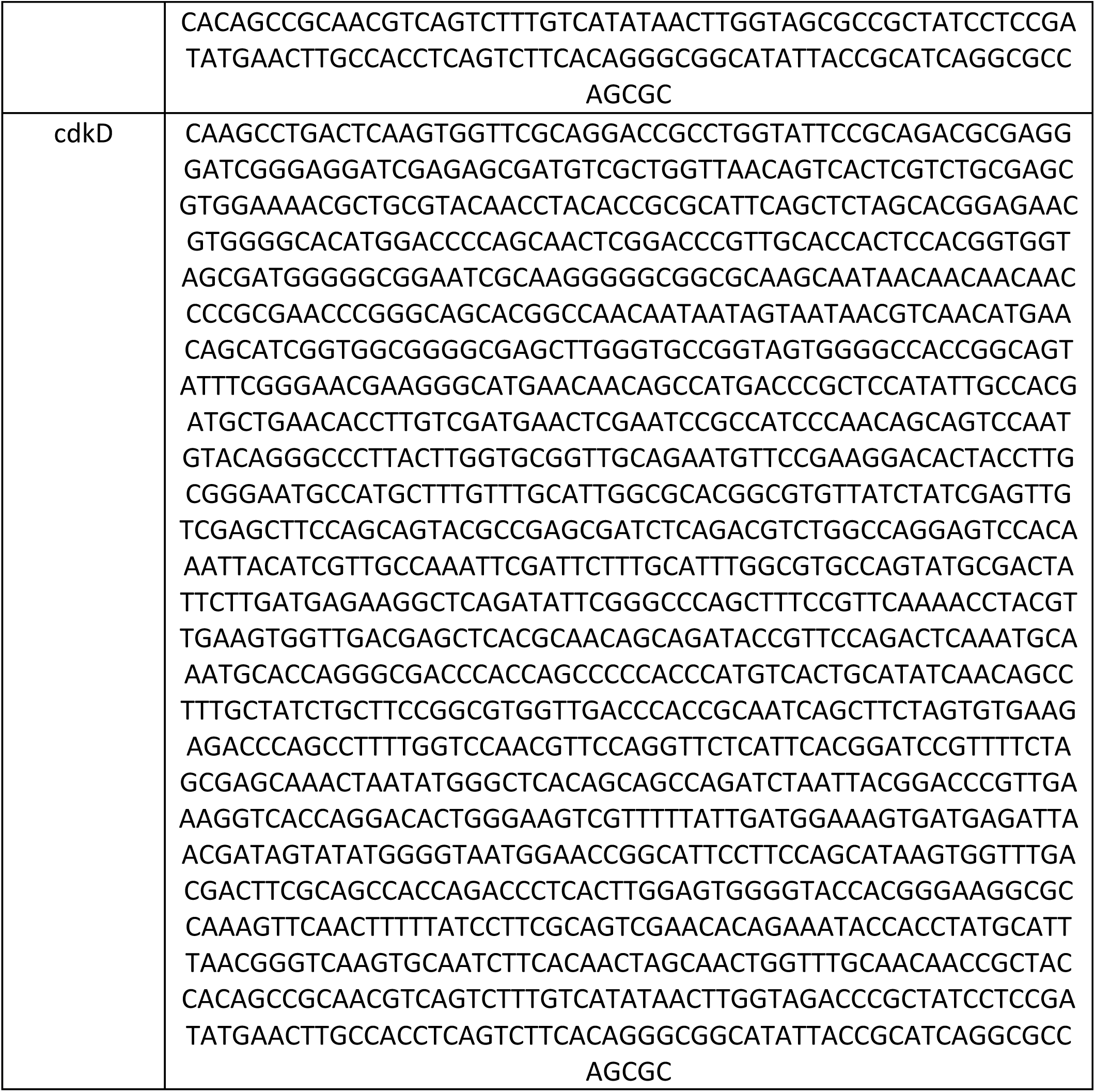
Geneblocks to create Whi3 mutants.

**Table 4.**
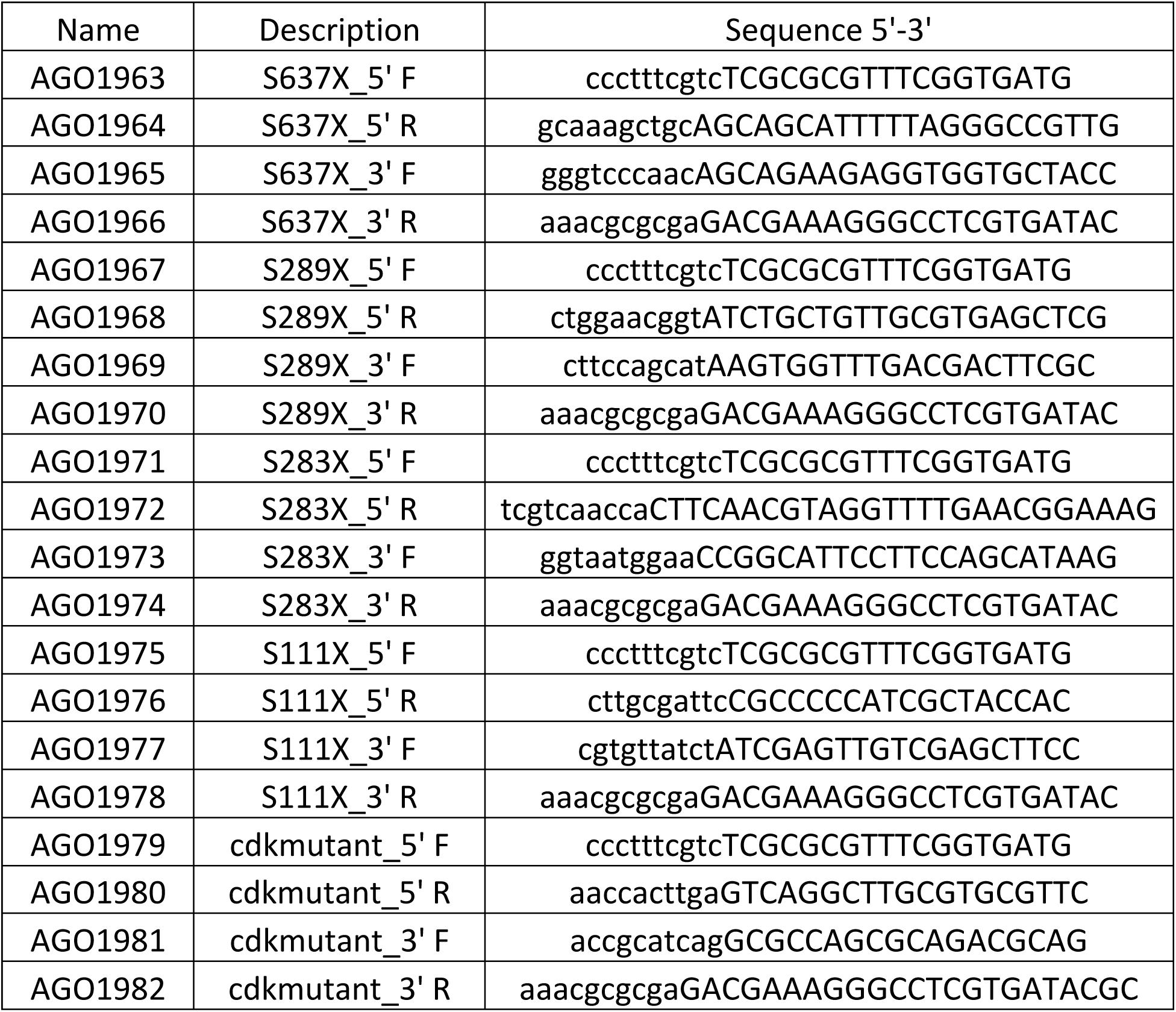
Primers to Create Gibson Fragments.

**Table 5.**
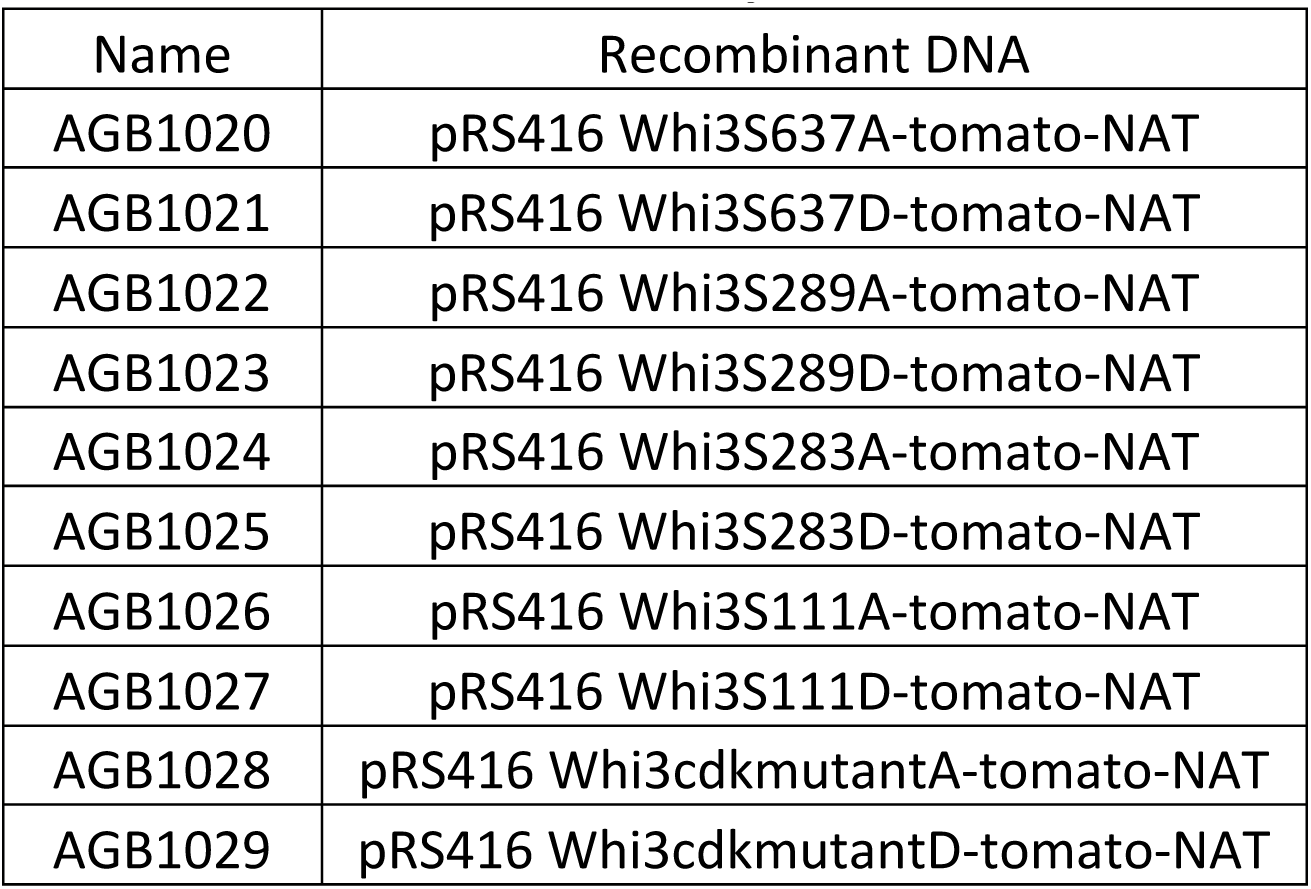
Plasmids created for this study.

| Name | Recombinant DNA |
| --- | --- |
| AGB1020 | pRS416 Whi3S637A-tomato-NAT |
| AGB1021 | pRS416 Whi3S637D-tomato-NAT |
| AGB1022 | pRS416 Whi3S289A-tomato-NAT |
| AGB1023 | pRS416 Whi3S289D-tomato-NAT |
| AGB1024 | pRS416 Whi3S283A-tomato-NAT |
| AGB1025 | pRS416 Whi3S283D-tomato-NAT |
| AGB1026 | pRS416 Whi3S111A-tomato-NAT |
| AGB1027 | pRS416 Whi3S111D-tomato-NAT |
| AGB1028 | pRS416 Whi3cdkmutantA-tomato-NAT |
| AGB1029 | pRS416 Whi3cdkmutantD-tomato-NAT |

**Table 6.** Strains used in this study.

| Name | Genotype |
| --- | --- |
| AG834 | $\Delta l\Delta t$-Whi3-tomato::NAT |
| AG853 | $\Delta l\Delta t$-Whi3(S637A)-tomato::NAT |
| AG854 | $\Delta l\Delta t$-Whi3(S637D)-tomato::NAT |
| AG855 | $\Delta l\Delta t$-Whi3(S289A)-tomato::NAT |
| AG856 | $\Delta l\Delta t$-Whi3(S289D)-tomato::NAT |
| AG857 | $\Delta l\Delta t$-Whi3(S283A)-tomato::NAT |
| AG858 | $\Delta l\Delta t$-Whi3(S283D)-tomato::NAT |
| AG859 | $\Delta l\Delta t$-Whi3(S111A)-tomato::NAT |
| AG860 | $\Delta l\Delta t$-Whi3(S111D)-tomato::NAT |
| AG861 | $\Delta l\Delta t$-Whi3(cdkmutA)-tomato::NAT |
| AG862 | $\Delta l\Delta t$-Whi3(cdkmutD)-tomato::NAT |

### Whi3 protein imaging and Analysis

*A. gossypii* cells were grown for 12.5h shaking at 150 rpm in Ashbya full media (AFM) at 30°C with appropriate selection, collected by centrifugation for 2 min at 500 rpm, and resuspended in 2X low fluorescence minimal media (LFM). Heat stressed cells were shifted to 42°C at 10.5h. Cells were arrested with 7.5 μg/mL nocodazole at 10.5h. Cells were mounted on 1.4% agarose gel pads made with LFM, sealed with VALAP, and then imaged using a widefield microscope (Nikon Eclipse TI stage) with a Plan Apo λ 60x/1.40 Oil Ph3 DM objective and an Andor Zyla 4.2 plus VSC-06258 camera. Images were deconvolved using 12 iterations of the Lucy-Richardson algorithm in Nikon Elements software and then processed for display using Fiji.

To measure Whi3 abundance, hyphal segments were traced in the phase channel in a middle plane to create ROIs. Z-series were used to create a sum intensity projection of the fluorescent channel, and then the mean fluorescent signal in each ROI was measured. These values were then averaged to generate Whi3 abundance in each strain. Abundances of mutant strains were then normalized to the control abundance. To measure hyphal tips containing Whi3 puncta, hyphal tips were marked in phase channel and then manually scored for the presence or absence of a Whi3 punctum in fluorescent channel. Mutant strains were compared to control using an N-1 Chi-Square test. To measure average number of Whi3 puncta in hyphae, 10 μm hyphal segments were traced in phase channel and then the number of Whi3 puncta in each ROI was counted manually. Mutant strains were compared to control using a T-test.

### Mass Spectrometry

For mass spectrometry, *A. gossypii* cells were grown for 16.5h shaking in Ashbya full media (AFM) at 30°C with appropriate selection. Heat stressed cells were shifted to 42°C at 14.5h. Cells were arrested with 7.5 μg/mL nocodazole at 14.5h. Cells were collected via vacuum filtration and rinsed with PBS before being resuspended in 2X-NP-40 buffer (6 mM Na2HPO4, 4 mM NaH2PO4xH2O, 1% NONIDET P-40, 150 mM NaCl, 2 mM EDTA, 50 mM NaF, 4 μg/ml leupeptin, 0.1 mM Na3VO4 + fresh 1X EDTA-free protease inhibitor, 1 mM PMSF, 0.01 mg/ml leupeptin, 50 mM β-glycerophosphate). Resuspended cells were flash frozen in liquid nitrogen, then lysed using a coffee grinder at 4°C. Lysate was clarified by spinning in a tabletop centrifuge at 13.K rpm and 4°C for 20 minutes. Supernatant was combined with IgG Sepharose beads and allowed to bind rotating at 4°C for 2h. Beads were washed with IPP 150 buffer (10 mM Tris-HCl pH 8.0, 150 mM NaCl, 0.1% NP-40) and resuspended in Denaturing Buffer (2% SDS, 50 mM Tris pH 8.1, 50 mM NaCl, 20% glycerol + fresh 2 mM DTT). Sample was incubated at 65°C for 20 minutes, then cooled to room temperature before adding 6mM iodoacetamide and vortexed. The sample was incubated at room temperature in the dark for 1 hour. DTT was then added to a final concentration of 5mM to quench. Samples were run in SDS-PAGE and bands corresponding to Whi3-TAP were excised and sent for mass spec.

Two independent experiments were sent for mass spec for each condition. Coverage of Whi3 in the first round: Asynchronous: 51.3%, Nocodazole Arrested: 52.1%, Heat Stress: 7.1%. Coverage of Whi3 in the second round: Asynchronous: 70.9%, Nocodazole Arrested: 73.9%, Heat Stress: 75.7%.

### Synchrony Imaging and Index

*A*.*gossypii* cells were grown for 16h shaking in Ashbya full media (AFM) at 30°C with appropriate selection then fixed with 3.7% formaldehyde shaking at 30°C for 1 hr. Cells were washed with PBS and resuspended in Solution A (100 mM K2PHO4 pH 7.5, 1.2M Sorbitol) and digested with zymolyase at 37°C to remove cell wall. Cells were washed with Solution A and then spotted onto polylysine treated wells on a slide. Cells were washed with PBS, blocked with BSA, and then incubated overnight with rat α-alpha tubulin antibody at 4°C. The following day cells were washed and incubated with fluorescently labeled secondary antibody + Hoechst. Cells were washed a final time then covered with Prolong gold mounting medium for imaging. Cells were imaged using a widefield microscope (Nikon Eclipse TI stage) with a Plan Apo λ 100x/1.45 Oil Ph3 DM objective and an Andor Zyla VSC-06258 camera. Images were deconvolved using 25 iterations of the Lucy-Richardson algorithm in Nikon Elements software and then processed using Fiji. Cell synchrony index was performed as described previously (Nair et al., 2010).

### Actin Zone Length Imaging and Analysis

*A. gossypii* cells were grown for 15h shaking in Ashbya full media (AFM) at 30°C with appropriate selection then fixed with 3.7% formaldehyde shaking at 30°C for 1 hr. Cells were collected by centrifugation, washed with PBS, then incubated in PBS with Alexa Fluor Phalloidin 488 at 6.6 μM for 1 hr. Cells were washed with PBS and mounted on glass slides with Prolong Gold mounting medium. Cells were imaged on a Zeiss LSM 880 AIRYSCAN with a Plan-Apochromat 63x/1.4 Oil DIC M27 objective. Images were airyscan processed using Zen software. Actin zone lengths were measured using Fiji. Mutant strains were compared to control using a T-test.

### Cell Branching Imaging

Ashbya spores were spotted onto agar plates made with AFM and appropriate selection. Cells were grown at 30°C and imaged on a Nikon Eclipse TS100 with a 10x/0.25 Ph1 ADL objective using an iDu Optics LabCam Microscope Adaptor and an iPhone 6 in LapseIt software.

---

**Supplemental Figure 1.**
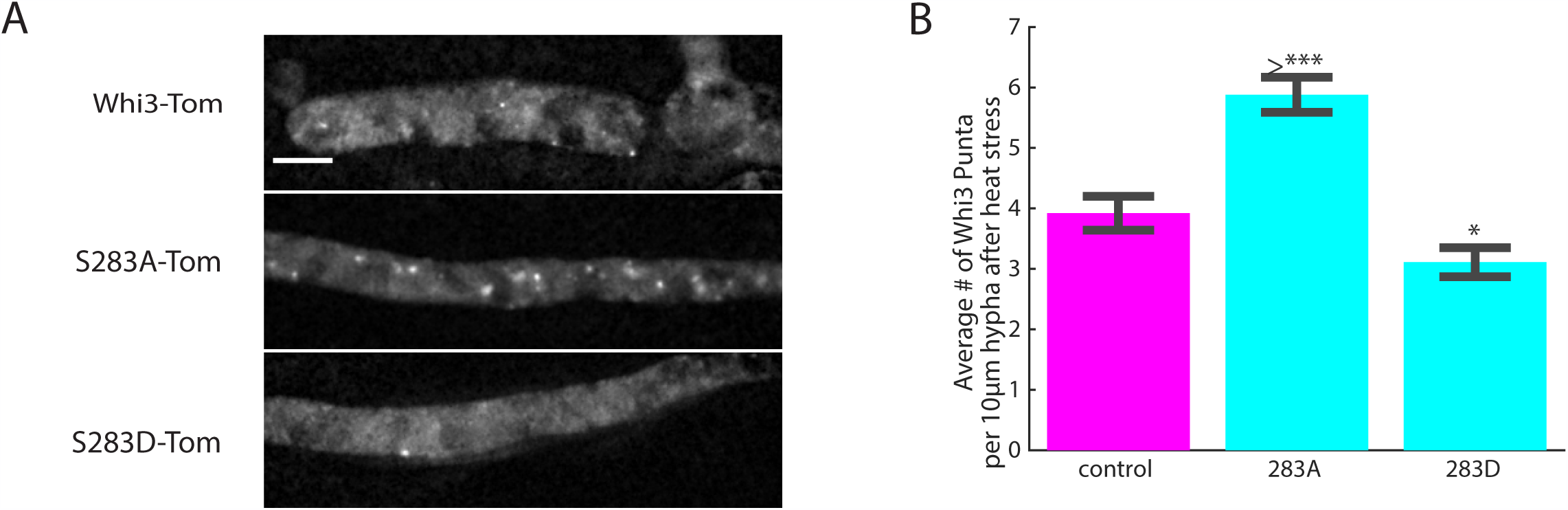
Residue S283 regulates Whi3 response to heat stress. (A) Representative images of Whi3 protein in control, 283A, and 283D strains localized in the hypha following heat stress. Images are maximum intensity projections. Scale bar 5 μm. (B) Average number of Whi3 puncta per 10 μm hyphal segment in control, 283A, and 283D strains localized in the hypha following heat stress. Bars denote SE. \**p* < 0.05 by T-test.

